# Metagenomic binning through low density hashing

**DOI:** 10.1101/133116

**Authors:** Yunan Luo, Y. William Yu, Jianyang Zeng, Bonnie Berger, Jian Peng

## Abstract

Bacterial microbiomes of incredible complexity are found throughout the world, from exotic marine locations to the soil in our yards to within our very guts. With recent advances in Next-Generation Sequencing (NGS) technologies, we have vastly greater quantities of microbial genome data, but the nature of environmental samples is such that DNA from different species are mixed together. Here, we present Opal for metagenomic binning, the task of identifying the origin species of DNA sequencing reads. Our Opal method introduces low-density, even-coverage hashing to bioinformatics applications, enabling quick and accurate metagenomic binning. Our tool is up to two orders of magnitude faster than leading alignment-based methods at similar or improved accuracy, allowing computational tractability on large metagenomic datasets. Moreover, on public benchmarks, Opal is substantially more accurate than both alignment-based and alignment-free methods (e.g. on SimHC20.500, Opal achieves 95% F1-score while Kraken and CLARK achieve just 91% and 88%, respectively); this improvement is likely due to the fact that the latter methods cannot handle computationally-costly long-range dependencies, which our even-coverage, low-density fingerprints resolve. Notably, capturing these long-range dependencies drastically improves Opal’s ability to detect unknown species that share a genus or phylum with known bacteria. Additionally, the family of hash functions Opal uses can be generalized to other sequence analysis tasks that rely on k-mer based methods to encode long-range dependencies.

---

Metagenomics is the study of the microbiome— the many genomes (bacterial, fungal, and even viral) that make up a particular environment. The microbiome has already been linked to human health: soil from a particular region can lead to the discovery of new antibiotics [1]; the human gut microbiome has been linked to Crohn's Disease [2], obesity [3] and even Autism Spectrum Disorder [4]. Metagenomics fundamentally asks what organisms are present in a genomic sample with the goal of gaining insight into function. However, the sequencing datasets required to shine any light on these questions are gigantic and vastly more complex than standard genomic datasets. This data results in major identification challenges for certain bacterial, as well as viral, species, strains, and genera [5, 6].

We focus on whole-genome metagenomic DNA sequencing, since cheaper Amplicon-based sequencing methods, which concentrate on the diversity of given marker genes (e.g. the 16S rRNA gene) and only analyze protein-coding regions, are limited in their ability to provide microbial functions from the samples [7, 8, 9]. Unfortunately, metagenomic sequencing data is inherently complex; the mixing of DNA from many different, sometimes related organisms in varying quantities poses substantial computational and statistical challenges to metagenomic binning, the process of grouping reads and assigning them to an origin organism. This important first step occurs before downstream data analysis can be applied to elucidate the structure of microbial populations and assign functional annotations [7]. Existing sequence alignment tools, such as BWA [10], Bowtie 2 [11] or BLAST [8], can readily be used and usually provide high-resolution alignments and accurate results by simply finding the highest scoring matching genome; they have the added advantage of tolerance to small numbers of mismatches or gaps. However, the computational cost of alignment-based methods becomes prohibitive as metagenomic datasets continue to grow [12, 13].

Alternatively, the field has turned to alignment-free metagenomic binning (also known as compositional binning) [14], which assigns sequence fragments to their taxonomic origins according to specific patterns of their constituent k-mers. State-of-the-art tools Kraken [12] and CLARK [15] use exact occurrences of uniquely discriminating k-mers in reads and are very efficient, but are limited in both their sensitivity and ability to detect unknown organisms. Other approaches rely on supervised machine learning (ML) classifiers, such as Naive Bayes or support vector machines (SVMs), trained on a set of reference genome sequences to classify the origins of metagenomic fragments [16, 17, 18, 19] using the relative k-mer frequency vector of a read. More recently, latent strain analysis performs covariance analysis of k-mers to partition reads for low-abundance strain assembly and detection [20]. All these approaches are often faster than alignment-based methods [10]. However, because they require exact matches of k-mers, these methods exhibit drawbacks including intolerance to mismatches or gaps; here we develop algorithmic tools to address these shortcomings.

As large k-mer sizes incur high memory usage and computing requirements (the space of k-mers grows exponentially in k), existing metagenomic binning methods generally work with a low fixed dimensionality (k): PhyloPythia [21] uses an ensemble of SVM models trained on contiguous 6-mers and its successor, PhyloPythiaS [17], further improves the binning accuracy by tweaking the SVM model and simultaneously including k-mers of multiple sizes (k = 3, 4, 5, 6) as compositional features. Some existing methods use mid-size k-mers (e.g. k=31), but primarily for fast indexing and nearest exact search [15, 22, 23, 12] and not in a supervised manner. Longer k-mers have the potential to capture compositional dependency within larger contexts because they span a larger section of the read. They can lead to higher binning accuracy but are also more prone to noise and errors if used in the supervised setting. To address this problem, locality-sensitive hashing (LSH) techniques, such as minHash [24] and randomly spaced k-mer construction have been developed for representing long k-mers sparsely [25], but as they are currently used in the high-density regime [22], they still run into the same exponential space problem of large k-mer sizes (Online Methods). However, to the best of our knowledge low-density hashing has not previously been used for metagenomic analysis.

Here we newly overcome these bottlenecks in handling long k-mers by developing a novel compositional metagenomic binning algorithm, Opal, which efficiently encodes long k-mers using low-dimensional profiles generated using even-coverage, low-density hashing. We take inspiration from low-density parity-check (LDPC) error correcting codes (also known as Gallager codes) to generate evenly-covering sets of random positions of a k-mer [26, 27], which we then apply to the machine learning pipeline introduced by Vervier, et al. [18] for metagenomic sequence classification. This innovation overcomes the limitations of uniformly random LSH functions, which despite their many nice theoretical properties, are typically not efficient for the task of constructing metagenomic fingerprints because of uneven coverage (Figure 1).

**Figure 1.**
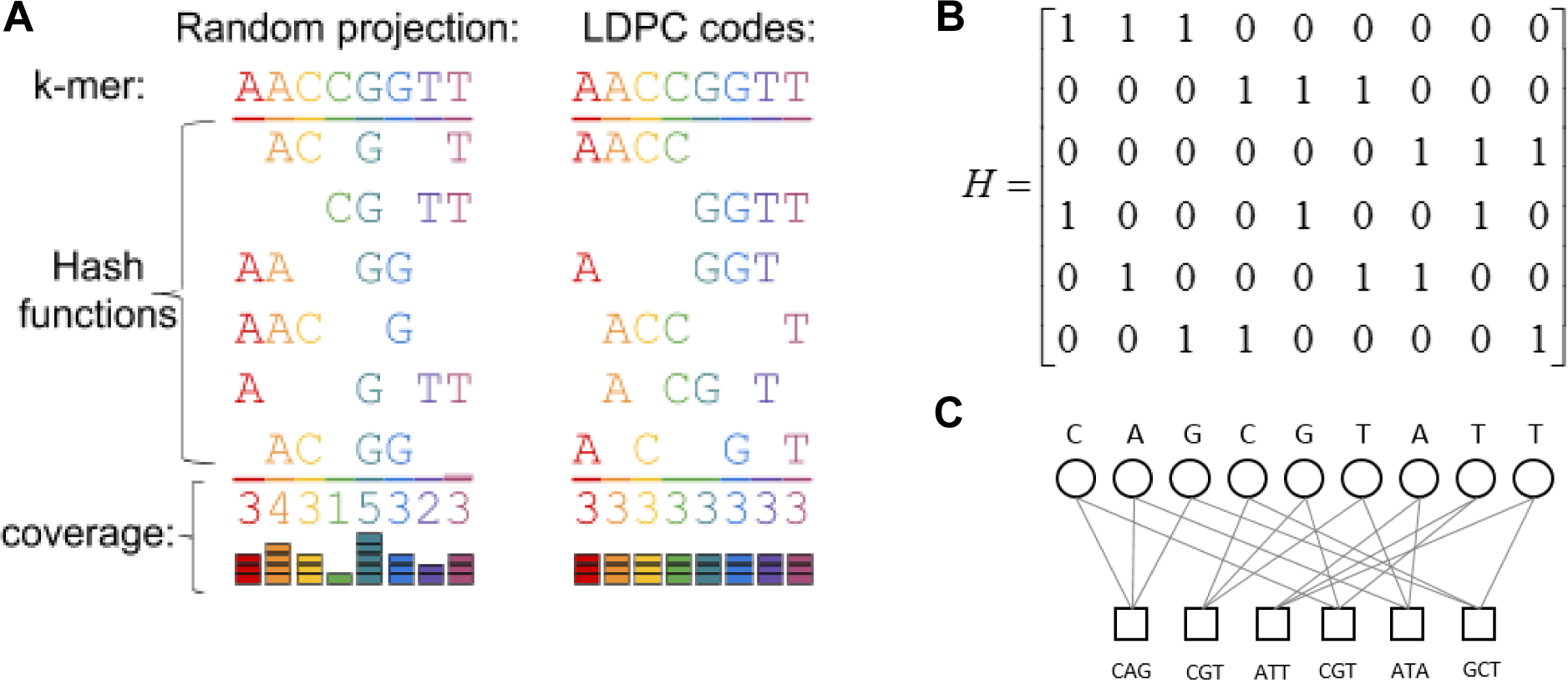
Low-density hashing with even coverage. ***(a)*** Random projections onto subspaces (left) cover all positions evenly only in expectation, and for small numbers of hash functions, will give uneven coverage. Using Gallager-inspired low density parity check (LDPC) codes allows us to guarantee even coverage of all positions in the k-mer (right) with a small number of hash functions. ***(b)*** Intuitively, one can think of a (k, t)-hash function as a 0/1 vector of length k with t 1’s specifying the locations in the k-mer that are selected. Given any (k, t)-hash function h (e.g. the vector with t 1’s followed by k-t 0’s), one can uniformly randomly construct another (k, t)-hash function by permuting the entries of h. The key to the Opal’s Gallager-inspired LSH design is that instead of starting with a single hash function and permuting it repeatedly, we start with a hash function matrix H which is a low-density parity check matrix. H is designed such that in the first row *h*_1_, the first t entries are 1, in the second row *h*_2_, the second t entries are 1, and so on, until each column of H has exactly one 1. Permuting the columns of H repeatedly generates random LSH functions that together cover all positions evenly, ensuring that we do not waste coding capacity on any particular position in the k-mer. Additionally, for very long k-mers, we can construct the Gallager LSH functions in a hierarchical way to further capture compositional dependencies from both local and global contexts (See *Online Methods*). ***(c)*** The rows of H are then used as hash functions.

Remarkably, when tested on a large dataset with 50 microbial species, Opal achieves both improved accuracy and up to two orders of magnitude improvement in binning speed on large datasets as compared to BWA-MEM [10], a state-of-the-art alignment-based method (Supplementary Fig S1-2); we can additionally use Opal as a first-pass coarse search [13, 28] before applying BWA-MEM for nearly 20 times speedup for the aligner (Supplementary Fig S2). As other compositional classifiers have similar speed gains over alignment-based methods, we shall henceforth focus on comparisons against compositional methods.

We offer two major conceptual advances in this work. First, although low-density LSH with uneven coverage has previously been used for fast sequence alignment and assembly [29, 9], it is the first time that it has been used for compositional metagenomic binning. Second, we have developed LSH functions based on the Gallager design for even coverage of very long k-mers (e.g. k = 64, 128), making the use of long k-mers practically possible. Of note, high density LSH (otherwise known as spaced-seeds) has been applied to metagenomic binning [2], but lowering the density is problematic without our second innovation to ensure even coverage of locations within a k-mer, as uneven coverage significantly decreases accuracy (Fig. 2). In this figure, we first importantly observe that low-density random long k-mer LSH provides better training accuracy than contiguous short k-mers [18], even when the feature space for the short k-mers is larger. Second, even coverage using Gallager codes provides another substantial decrease in the classification error; as substitution error rate increases, Opal’s advantages become ever more apparent.

**Figure 2.**
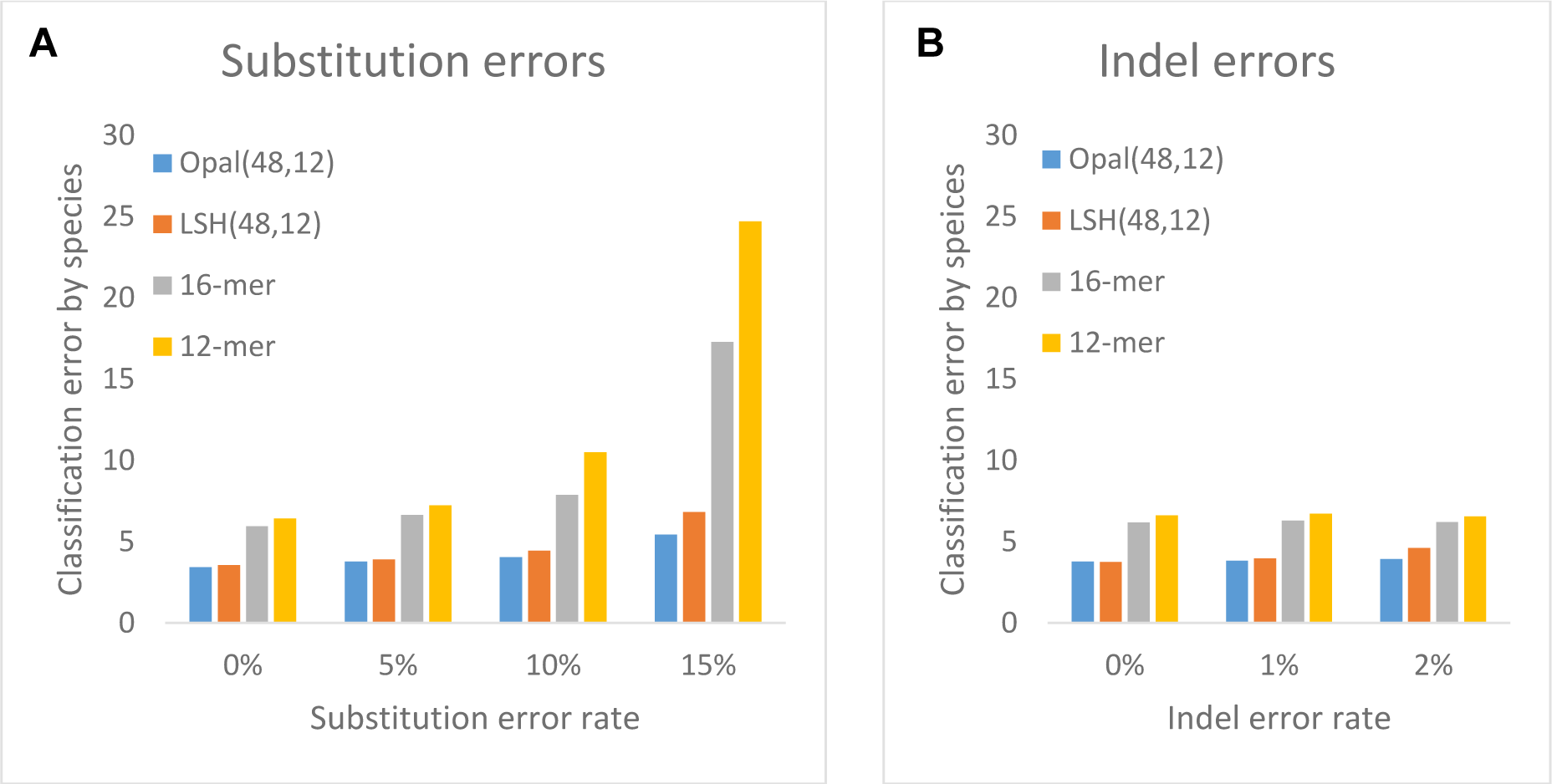
Comparison of Opal against compositional SVM-based approaches. On a synthetic dataset of fragments of length 200 drawn from an in-house dataset of 50 bacterial species, using Opal hash functions as features outperforms uniformly random locality sensitive hash (LSH) functions, as well as using contiguous 16-mers and 12-mers, with **(a) substitution errors** and **(b) indels**. We note particularly good robustness against substitution errors.

Opal outperforms the Kraken [12] and Clark [15] classifiers at assigning reads to both known species and to higher phylogenetic levels for unknown species (Figure 3). On three published benchmarks of real and simulated data with either 10 or 20 species used in previous testing of Kraken and Clark [12, 15], Opal outperforms both methods when trained on 24-mers with 2 hashes of row-weight 12 (Fig 3a). We also compared Opal to MetaPhlAn2 [30] for metagenomic profiling; even using their (MetaPhlAn2’s) marker genes, Opal performs better on the species and genus levels (Supplementary Table S3). *Opal thus achieves better accuracy than both alignment-based and existing compositional k-mer methods for classifying known species, at improved or similar runtimes.*

**Figure 3.**
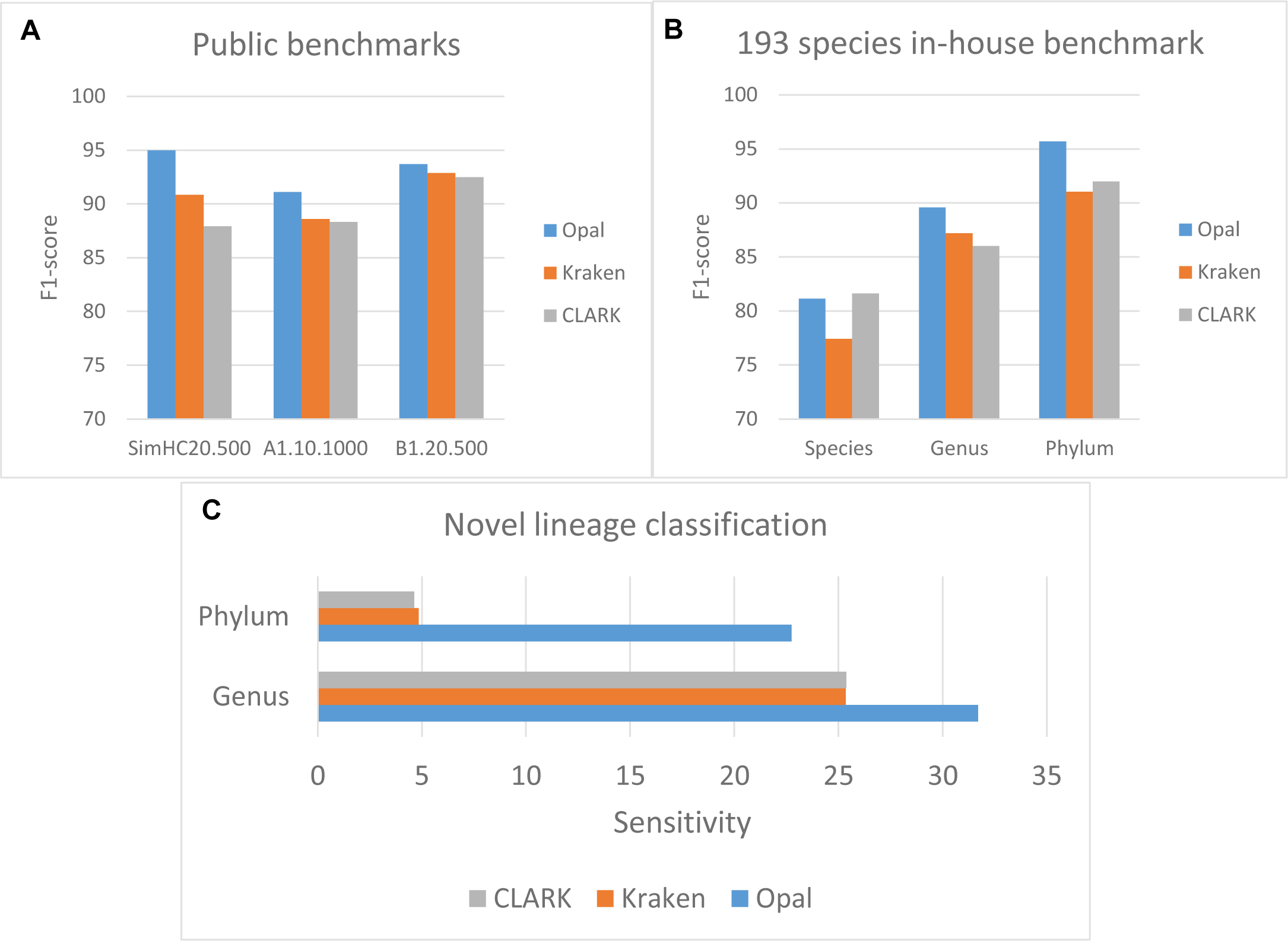
Comparison of Opal against Kraken and CLARK. ***(a)*** Opal achieves higher classification accuracies on three public benchmark data sets than two other state-of-the-art compositional classifiers. ***(b)*** Opal’s performance increase is especially pronounced at higher phylogenetic levels on a benchmark set of 193 species from the literature [18]; for the genus-level study, we trained Opal using the genus as the class label instead of the species, and similarly for the phylum-level study. ***(c)*** This increase allows Opal to have greater sensitivity to novel lineages, where the source genomes of the sequenced reads share either a genus or phylum, but not lower phylogenetic levels, with the training data Opal is given. That is, for the genus-level comparison, we removed a species from the dataset, and then trained at the genus-level on the remaining species, finally testing if we could correctly identify the genus of the removed species from its reads.

Notably, Opal’s performance increase is especially pronounced at higher phylogenetic levels (Fig 3b). When tested on a large benchmark of 193 species [18], Opal demonstrates greater sensitivity to novel lineages, where the source genomes of the sequenced reads share either a genus or phylum but are dissimilar at lower phylogenetic levels (Fig 3c). By detecting the genus or phylum of reads originating from unidentified species, Opal enables scientists to perform further analyses on reads by starting with information on the phylogenetic histories of those unknown species.

Additionally, Opal is effective at the subspecies level. When trained on subspecies references, even for seven closely related subspecies of *E. coli*, Opal disambiguates error-free synthetic reads with <15% classification error, while Kraken and CLARK both had over 30% classification error (Supplemental Fig S3). For subspecies classification, we found it necessary to train at a much higher depth of simulated read coverage than higher-order classification to increase accuracy to acceptable levels, likely due to the fact that related subspecies share many substrings of nucleotides in their genomes.

Not only is Opal a drop-in tool for metagenomic analysis pipelines (e.g. Vervier, et al. [18]), but the ideas that went into its construction can also potentially be applied to improve the discriminative power of other methods. The Opal Gallager LSH functions can immediately be used in lieu of contiguous k-mers in other metagenomic tools, such as Latent Strain Analysis [20]. Our method can also be seen as a new dimensionality reduction approach for genomic sequence data, extending the ordinary k-mer profile-based methods with compressed signatures, or fingerprints, of the reads.

With improvements in metagenomic sequencing technologies producing ever larger amounts of raw data, fast and accurate classifiers will become essential for handling the data deluge. Here we show that with a straightforward modification to the choice of hash functions, we can substantially improve feature selection and thus accuracy over other state-of-the-art classifiers. This improved accuracy manifests itself most strongly at higher phylogenetic levels, allowing Opal to better classify reads originating from unknown species. We expect Opal to be an essential component in the arsenal of metagenomic analysis toolkits.

The Opal software (available at http://opal.csail.mit.edu and https://github.com/yunwilliamyu/opal) will greatly benefit any researchers who are producing and analyzing large amounts of environmental metagenomic sequencing data.

## Acknowledgments

An extended abstract of an earlier version of this work appeared in RECOMB 2016. Y.W.Y and B.B. are partially supported by NIH grant GM108348 and an MIT Center for Microbiome Informatics and Therapeutics Pilot Grant (to B.B.). Y.W.Y gratefully acknowledges support from the Fannie and John Hertz Foundation. We thank Moran Yassour for introducing us to the subspecies classification problem and Ashwin Narayan for fruitful discussions.

## Author Contributions

J.P. and B.B. conceived the central premise of using low density locality sensitive hashing for metagenomic binning and guided the study. Initial development, coding, and all experimentation except for subspecies classification was performed by Y.L. and J.Z. with guidance from J.P. and B.B. Reframing the study, rewriting the code in Python, preparing a user-friendly UI, and running all subspecies classification experiments were done by Y.W.Y. Writing was a joint endeavor of all listed authors.

## Competing financial interests

The authors have no competing financial interests.

## Online Methods for

### Compositional read classification with *k*-mer profiles

We assume that a sequence fragment *s* ∈ *Σ*^*L*^, where *Σ* = {*A*, *T*, *G*, *C*}, contains *L* nucleotides. A *k*-mer, with *k* < *L*, is a short word of *k* contiguous nucleotides. We define the *k*-mer profile of *s* in a vector representation 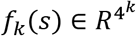. If we index each *k*-mer as a binary string with length 2*k*, then we have a one-to-one mapping between any *k*-mer and an integer from 0 to 2^2*k*^. In the rest of the paper, we will not distinguish the *k*-mer string with its integer presentation *i* for notational simplicity. Each coordinate in the *k*-mer profile *f*_*k*_(*s*,*i*) stores the frequency of *k*-mer *i* in the sequence fragment *s*. For instance, for a fragment *s* = *AATTAT*, its 2-mer profile *f*_2_(*s*) has 4 non-zero entries: *f*_2_(*s*, *AA*) = 1/5, *f*_2_(*s*, *TT*) = 1/5, *f*_2_(*s*, *AT*) = 2/5 and *f*_2_(*s*, *TA*) = 1/5. In this way, instead of representing a *L*-nucleotide fragment in *O*(4^*L*^), we can use *k*-mer profile to represent it in *O*(4^*k*^). Similarly, we can construct *k*-mer profiles given hash functions that specify other positional subsequences of the *k*-mer, rather than only contiguous subsequences.

After the *k*-mer profile has been constructed, we can use supervised machine learning classification algorithms, such as logistic regression, naive Bayes classifier and support vector machines, to train a binning model. The training data can be generated by sampling *L*-nucleotide fragments from the reference genomes with taxonomic annotations. In particular, in this paper, we used one-against-all support vector machines, implemented using Vowpal Wabbit. Further details are given for specific experiments.

### Locality sensitive hashing

LSH is a family of hash functions that have the property that two similar objects are mapped to the same hash value [31]. For the metagenomic binning problem, we are only interested in strings of length *k*.Then a family of LSH functions can be defined as functions *h*: Σ^*k*^ → *R*^*d*^ which map *k*-mers into a *d*-dimensional Euclidean space. Assume that we consider Hamming distances between *k*-mers, if we choose *h* randomly and for two *k*-mers *s*_1_ and *s*_2_ with at most *r* different positions, *h*(*s*_1_) = *h*(*s*_2_) holds with probability at least *P*_1_. For two *k*-mers *s*_3_ and *s*_4_ with more than *R* different positions, *h*(*s*_3_) ≠ *h*(*s*_3_) holds with probability at least *P*_2_. With the construction of a LSH family, we can amplify *P*_1_ or *P*_2_ by sampling multiple hash functions from the family. Compared with the straightforward k-mer indexing representation, the LSH scheme can be more compact and more robust. For example, we can construct LSH functions such that d ≪ 4*k*. Moreover, when a small number of sequencing errors or mutations appear in the *k*-mer, LSH can still map the noisy *k*-mer into a feature representation that is very similar to original *k*-mer. This observation is highly significant since mutations or sequencing errors are generally inevitable in the data, and we hope to develop compositional-based methods less sensitive to such noises. One way to construct LSH functions on strings under Hamming distance is to construct index functions by uniformly sampling a subset of positions from the k-mer. Specifically, given a string *s* of length *k* over Σ, we choose *t* indices *i*_1_, …, *i*_*t*_ uniformly at random from {1, …, *k*} without replacement. Then, the spaced (*k*, *t*)-mer can be generated according to *s* and these indices. More formally, we can define a random hash function *h*: Σ^*k*^ → Σ^*t*^ to generate a spaced (*k*, *t*)-mer explicitly:

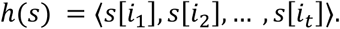

The hash value *h*(*s*) can also be seen as a 4^*t*^ dimensional binary vector with only the string *h*(*s*)’s corresponding coordinate set to 1 and otherwise 0. It is not hard to see that such LSH function *h* has the property that it maps two similar *k*-mers to the same hash value with high probability. For example, consider two similar *k*-mers *s*_1_ and *s*_2_ that differ by at most *r* nucleotides, then the probability that they are mapped to the same value is given by

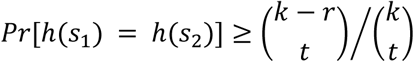

For two k-mers *s*_3_ and *s*_4_ that differ at least *R* nucleotides, the probability that they are mapped to different value is given by

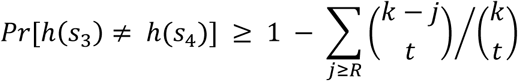

With the family of LSH functions, we randomly sample a set of *m* LSH functions and concatenate them together as the feature vector for a long *k*-mer. Note that the complexity of the LSH-based feature vector is only *O*(*m*4^*t*^), much smaller compared to *O*(4^*k*^) that is the complexity of the complete *k*-mer profile, so long as *t* is much smaller than *k*. As an aside, this is the reason that high-density hashing still runs into the exponential space blow-up problem. When *t* = *ck*, for some constant *c* > 0, *O*(4^*ck*^)is still exponential in *k*. It is for this reason that we turn to low-density hashing, where *t* s a small constant, in the next section.

More importantly, the LSH-based feature vector is not sensitive to substitution errors or mutations in the *k*-mer if *m* and *t* are well chosen, but for the traditional k-mer profile, even one nucleotide change can change the feature vector completely. To compute the feature vector for a metagenomic fragment with length *L*, we first extract all *k*-mers by sliding a window of length *k* over the sequence, and then apply *h* on each *k*-mer to generate LSH based feature vectors and then normalize the sum of the feature vectors by *L* – *k +* 1. In this way, one can easily show that similar fragments can also be mapped to similar LSH-based feature vectors. After the feature vectors are generated for fragments with taxonomic annotations, we train a linear classifier for metagenomic binning. It is also fairly straightforward to show that similar fragments have similar classification responses if the coefficients of the linear classification function are bounded. One may expect that the complexity of linear classification with k-mer profiles would be lower since there are at most *L* – *k +* 1 different *k*-mers in a fragment and can be computed easily using sparse vector multiplications, but we find that the LSH based feature vector is also sparse in practice and the indexing overhead is much smaller when constructing the feature vectors, since the LSH-based method can have much smaller dimensionality. In practice, the LSH-based methods can sometimes be even faster if *m* and *t* are not too large.

### Gallager low-density locality-sensitive hashing

Despite that the random LSH function family described above has a lot of nice theoretical properties, uniformly sampled LSH functions are usually not optimal in practice. Theoretical properties of LSH functions hold probabilistically, which means that we need to sample a large number of random LSH functions to make sure the bounds are tight. However, practically, we simply cannot use a very large number of random LSH functions to build feature vectors for metagenomic fragments, given the limited computational resources. Thus it would be ideal if we could construct a small number of random LSH functions that are sufficiently discriminative and informative to represent long *k*-mers. Here we take inspiration from the Gallager code or low-density parity-check code that has been widely used for noisy communication. The idea behind the Gallager code is similar to our LSH family but with a different purpose, namely error correction. The goal of the LDPC code is to generate a small number of extra bits when transmitting a binary string via a noisy channel [26, 27]. These extra bits are constructed to capture the long-range dependency in the binary string before the transmission. After the message string and these extra bits have been received, a decoder can perform error correction by performing probabilistic inference to compare the differences between the message string and these code bits to infer the correct message string. In the same spirit, we here adopt the idea behind the design of the LDPC code to construct a compact set of LSH functions for metagenomic binning.

To construct compact LSH functions, we hope to not waste coding capacity on any particular position in the *k*-mer. While, under expectation, uniformly sampled spaced (*k, t*)-mers on average cover each position equally, with a small number of random LSH functions, it is likely that we will see imbalanced coverage among positions since the probability of a position being chosen is binomially distributed. The Gallager’s design of LDPC, on the other hand, generates a subset of positions not uniformly random but make sure to equally cover each position [26]. So we can use the Gallager’s design to generate spaced (*k, t*)-mers. The Gallager’s LDPC matrix *H* is a binary matrix with dimension *m* × *k*, and has exactly *t* 1’s in each rows and *w* 1’s in each column. The matrix *H* can be divided into *w* blocks with *m*/*w* rows in each block. We first define the first block of rows as an 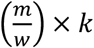 matrix *Q*:

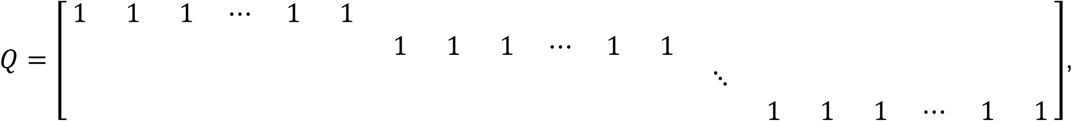

where each row of matrix Q has exactly t consecutive 1’s from left to right across the columns. Every other block of rows is a random column permutation of the first set, and the LDPC matrix H is given by:

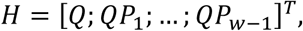

where *P*_*i*_ is a uniform random *n* × *n* permutation matrix for *i* = 1, …, *w* – 1. An example with *k* = 9, *t* = 3, *m* = 6, *w* = 2 is shown in **Figure 1**. An equivalent bipartite graph with the Gallager design matrix as the adjacency matrix also is shown. The algorithm for constructing the LDPC design matrix is as follows:

Algorithm 1: Gallager’s LDPC Matrix:

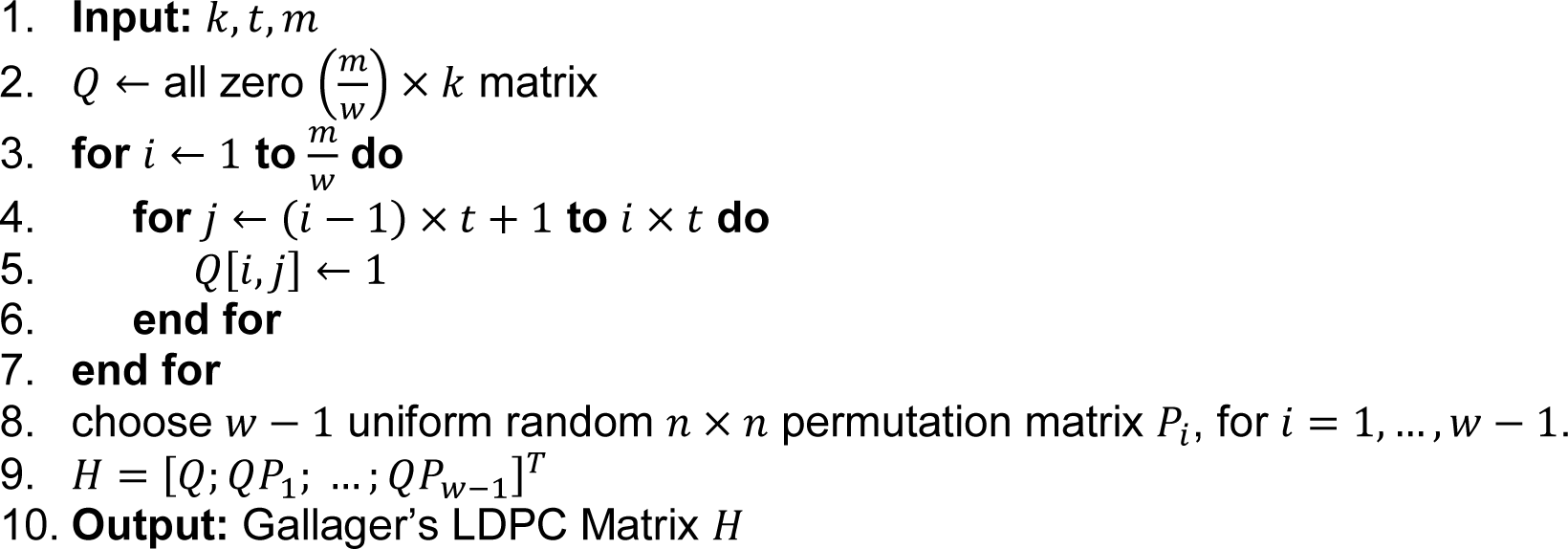

We use each row of *H* to extract a spaced (*k*, *t*)-mer to construct an LSH function. Note that the first set of *H* gives contiguous *t*-mers. With *m* Gallager LSH functions, we can see that each position in a *k*-mer is equally covered *w* times, while the same *m* uniformly sampled LSH function is very likely to have very imbalanced coverage times for different positions because of the high variance 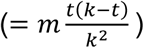. To further improve the efficiency, we construct random LSH functions with minimal overlap using a modified Gallager design algorithm. The idea is to avoid the “4-cycles” in the bipartite graph representation, as we hope not to encode two positions together in two “redundant” LSH functions [27]. An algorithm which finds “4-cycles” and removes them is shown here:

Algorithm 2: Removing 4-cycles:

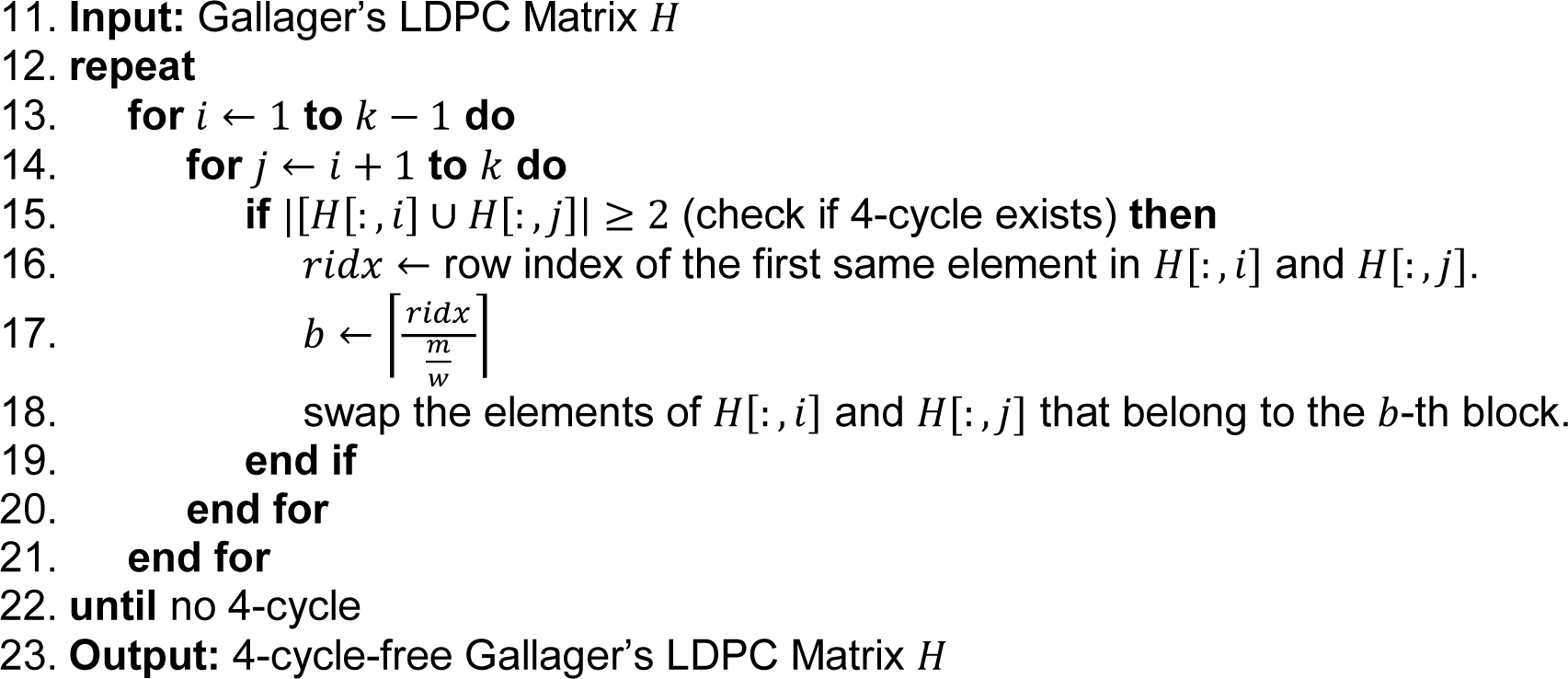

For very long *k*-mers, we can use a hierarchical approach to generate low-dimensional LSH functions for very long-range compositional dependency in *k*-mers. We first generate a number of intermediate spaced (*k*, *l*)-mers using the Gallager’s design matrix. Then from these (*k*, *l*)-mers, we again apply the Gallager’s design to generate (*l*, *t*)-mers to construct the (*k*, *l*, *t*) hierarchical LSH functions.

## Benchmarks

### Comparisons against Vervier, et al. SVM approaches (Figure 2)

For the synthetic benchmark we used in measuring the robustness of using Opal’s evenly spaced hashes for SVM features (**Figure 2**), we started with 50 full bacterial genomes in Fasta format downloaded from NCBI database.

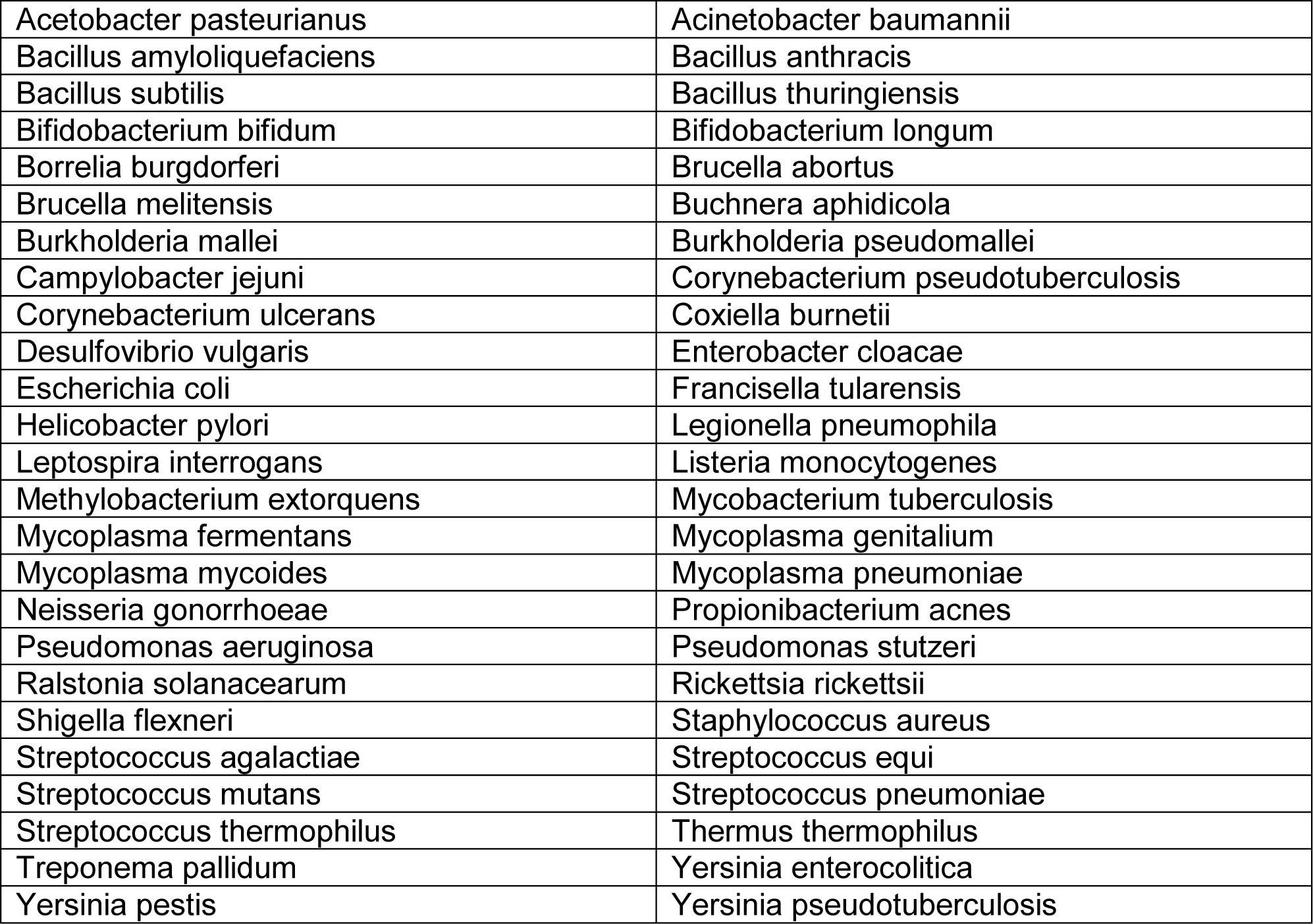

For training the SVM methods, synthetic reads of length 200 bp were randomly drawn from the bacterial genomes such that average depth of coverage was 5x; features from these reads were passed to Vowpal Wabbit 8.1.1 following the method of Vervier, et al.

Matching the behavior of Vervier, et al., we trained on 12-mer and 16-mer features. Additionally, Opal and random LSH features were chosen by taking 12 locations in k-mers of size 48. For the substitution error experiments, Opal and random LSH used 8 hash functions in addition to a contiguous 12-mer as features. For the indel error experiments, Opal and random LSH used 16 hash functions in addition to a contiguous 12-mer as features.

For testing, synthetic reads of of length 200 bp were again randomly drawn, but with average depth of coverage only 1x. For substitution error benchmarks, for each location in a read, with probability 0.05, 0.10, or 0.15, we replaced it uniformly randomly with one of the four nucleotides (i.e. one quarter of the time, despite a location being selected for a substitution error, it remained unchanged). For indel error benchmarks with indel error rates of 0.01 or 0.02, for each read, [read-length=200] * [indel rate] locations were selected to be indels. With equal probability, either that location is deleted, or a random base is inserted.

Classification error by species was computed by getting the classification error of reads from each species separately, and then averaging over all 50 species.

### Comparisons against Kraken and CLARK (Figure 3)

We compared the performance of Opal against Kraken and CLARK on the public benchmark datasets SimHC20.500, A1.10.1000, and B1.20.500 of 20, 10, and 20 species respectively.

The SimHC20.500 synthetic read dataset was previously used as a benchmark in the paper introducing CLARK [15], containing the following species:

|  |  |
| --- | --- |
| Alkaliphilus metalliredigens | Bradyrhizobium sp. BTAi1 |
| Burkholderia ambifaria | Chelativorans sp. BNC1 |
| Clostridium thermocellum | Dechloromonas aromatic |
| Desulfitobacterium hafniense | Frankia sp. Ccl3 |
| Geobacter metallireducens | Marinobacter aquaeolei |
| Methanosarcina barkeri | Nitrobacter hamburgensis |
| Nocardioides sp. JS614 | Polaromonas sp. JS666 |
| Pseudoalteromonas atlantica | Pseudomonas fluorescens |
| Rhodobacter sphaeroides | Rhodopseudomonas palustris |
| Shewanella sp. MR-7 | Syntrophobacter fumaroxidans |

The A1.10.1000 dataset of real sequencing was previously used as a benchmark in [32], containing the following species:

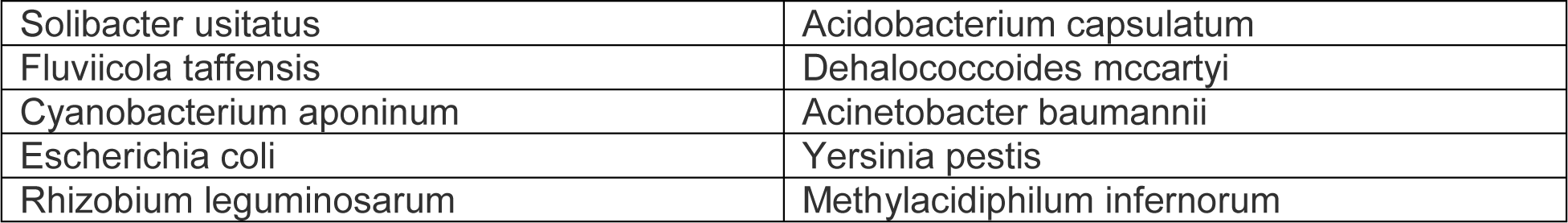

The B1.20.500 dataset of real sequencing reads was previously used as a benchmark in [32], containing the following species:

|  |  |
| --- | --- |
| Terriglobus saanensis | Propionibacterium avidum |
| Prevotella melaninogenica | Cyclobacterium marinum |
| Roseiflexus sp. RS-1 | Desulfurococcus kamchatkensis |
| Geitlerinema sp. PCC 7407 | Stanieria cyanosphaera |
| Thermococcus onnurineus | Bacillus licheniformis |
| Geobacillus sp. Y4.1MC1 | Roseburia hominis |
| Rickettsia Canadensis | Yersinia pestis |
| Ehrlichia ruminantium | Pantoea annatis |
| Zymomonas mobilis | Streptococcus dysgalactiae |
| Rhizobium leguminosarum | Akkermansia muciniphila |

Kraken was run with default options. CLARK was run with default options with k=31. Opal was run with L=100, depth-of-coverage=5, k=24, t=12, and 2 hashes (one contiguous and one spaced).

We additionally compared the performance of all three tools on a large 193 species benchmark used in [18] (the “medium” dataset in the referenced paper), containing the following species (listed here only by NCBI taxonomic ID for the sake of brevity):

> 24, 139, 154, 160, 172, 173, 174, 195, 196, 197, 210, 213, 235, 263, 274, 287, 294, 300, 303, 316, 339, 346, 347, 380, 382, 384, 408, 470, 485, 487, 518, 519, 520, 548, 550, 552, 553, 554, 571, 573, 615, 621, 622, 623, 624, 630, 632, 633, 636, 644, 666, 670, 672, 715, 727, 738, 770, 779, 782, 783, 785, 788, 803, 813, 817, 876, 881, 920, 948, 1085, 1096, 1245, 1280, 1282, 1304, 1307, 1308, 1309, 1311, 1313, 1314, 1318, 1328, 1334, 1336, 1338, 1351, 1352, 1390, 1392, 1396, 1398, 1402, 1404, 1406, 1423, 1428, 1488, 1491, 1502, 1513, 1515, 1534, 1579, 1580, 1581, 1582, 1584, 1587, 1590, 1598, 1604, 1613, 1624, 1639, 1681, 1685, 1717, 1718, 1719, 1747, 1764, 1765, 1767, 1769, 1772, 1773, 1804, 1833, 1912, 2096, 2102, 2105, 2115, 2209, 2261, 2285, 2287, 2743, 13373, 28025, 28035, 28197, 28450, 29449, 29459, 29461, 29501, 32046, 33959, 33990, 34021, 35554, 35791, 35794, 36809, 36855, 39152, 39491, 39492, 43080, 43771, 47715, 49338, 52584, 53399, 55601, 57975, 61624, 62322, 65058, 76759, 76860, 77038, 78331, 79967, 82996, 83554, 83558, 85991, 95486, 106590, 120577, 138563, 152480, 155892, 161493, 191026, 216816, 283734, 315405, 380021, 657445

Opal was trained at three different taxonomic levels of species, genus, and phylum, separately, with the same options as above. Kraken and CLARK are taxonomically aware, and automatically also attempt to give genus and phylum information.

For novel lineage classification, we removed a species/genus from the training data, but kept it in the test set, and then measured the sensitivity of classifying those reads at the higher genus/phylum level.

### Data Availability

All data used in this paper has been previously published and can be accessed through the references given above.

### Code Availability

Source code for Opal can be found online at http://opal.csail.mit.edu, and through the linked Github repository. All code has been published under the GNU General Public License.

## Supplementary Information

**Figure S1.**
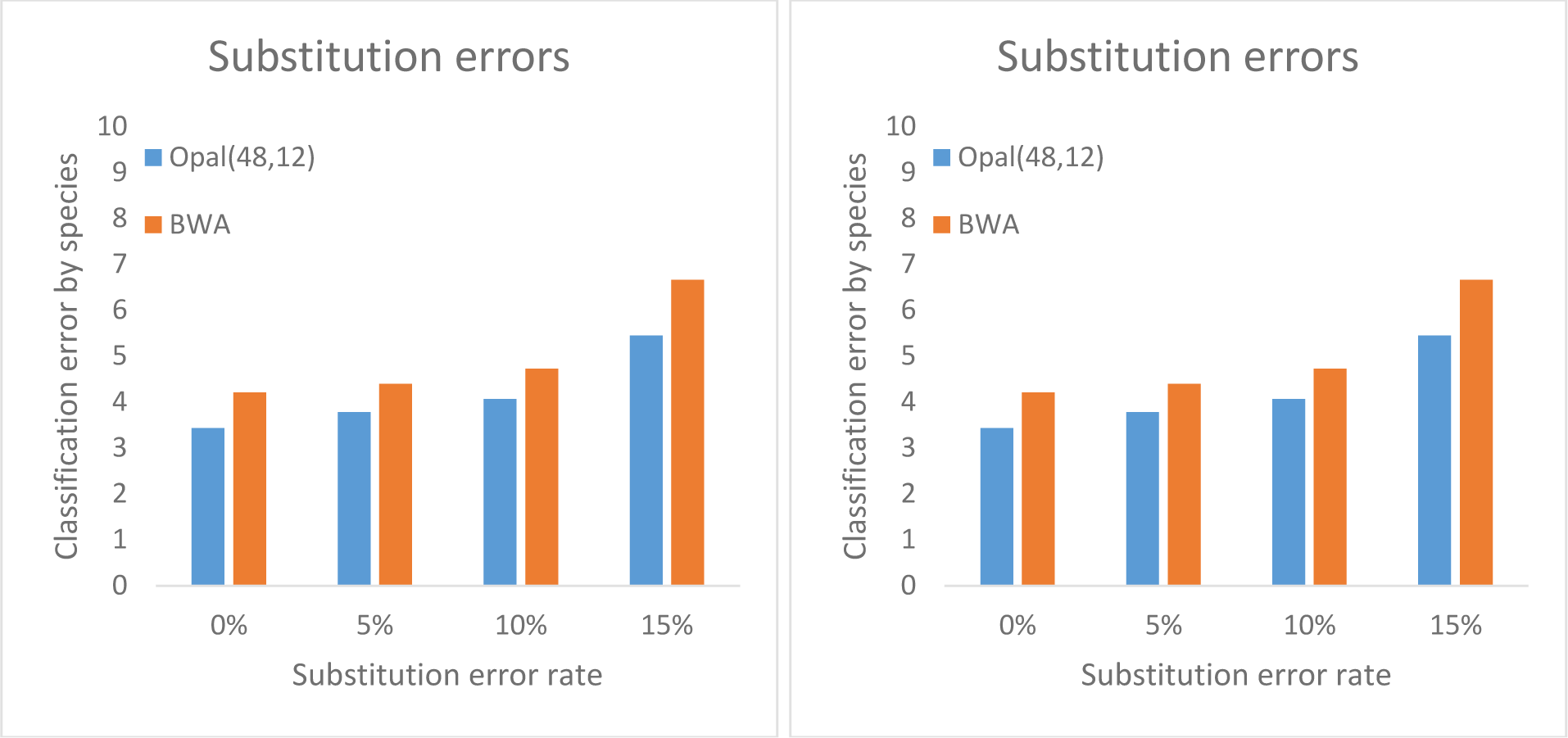
Opal is more accurate than BWA-MEM for classification. On the same synthetic benchmark used in **Figure 2** of fragments of length 200 drawn from an in-house dataset of 50 bacterial species, using Opal hash functions as features outperforms BWA-MEM for classification. Again, we not particular robustness against substitution errors.

**Figure S2.**
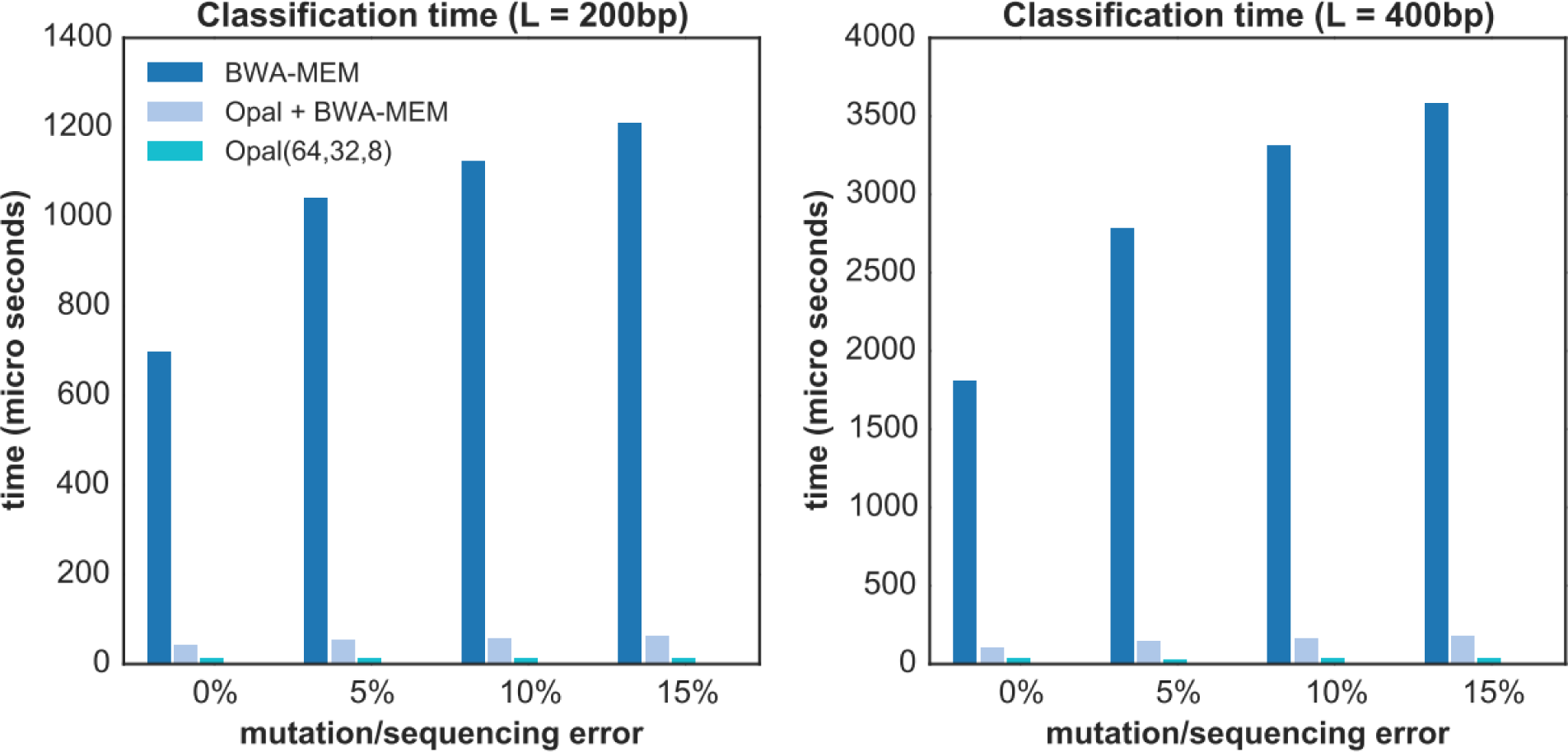
Opal run alone with parameters (64,32,8) achieves two orders of magnitude speedup over a state-of-the-part alignment-based method, BWA-MEM, on fragments of length (L) 200 and 400 at varying mutation/sequencing error rates (lower is better). We observe that compositional-based binning as “coarse search” for alignment-based methods can significantly speed up alignment time (Opal + BWA-MEM). In particular, Opal applied as a “coarse-search” procedure reduces the taxonomic space for a subsequent alignment-based BWA-MEM “fine search” to achieve nearly 20 times speedup.

**Figure S3.**
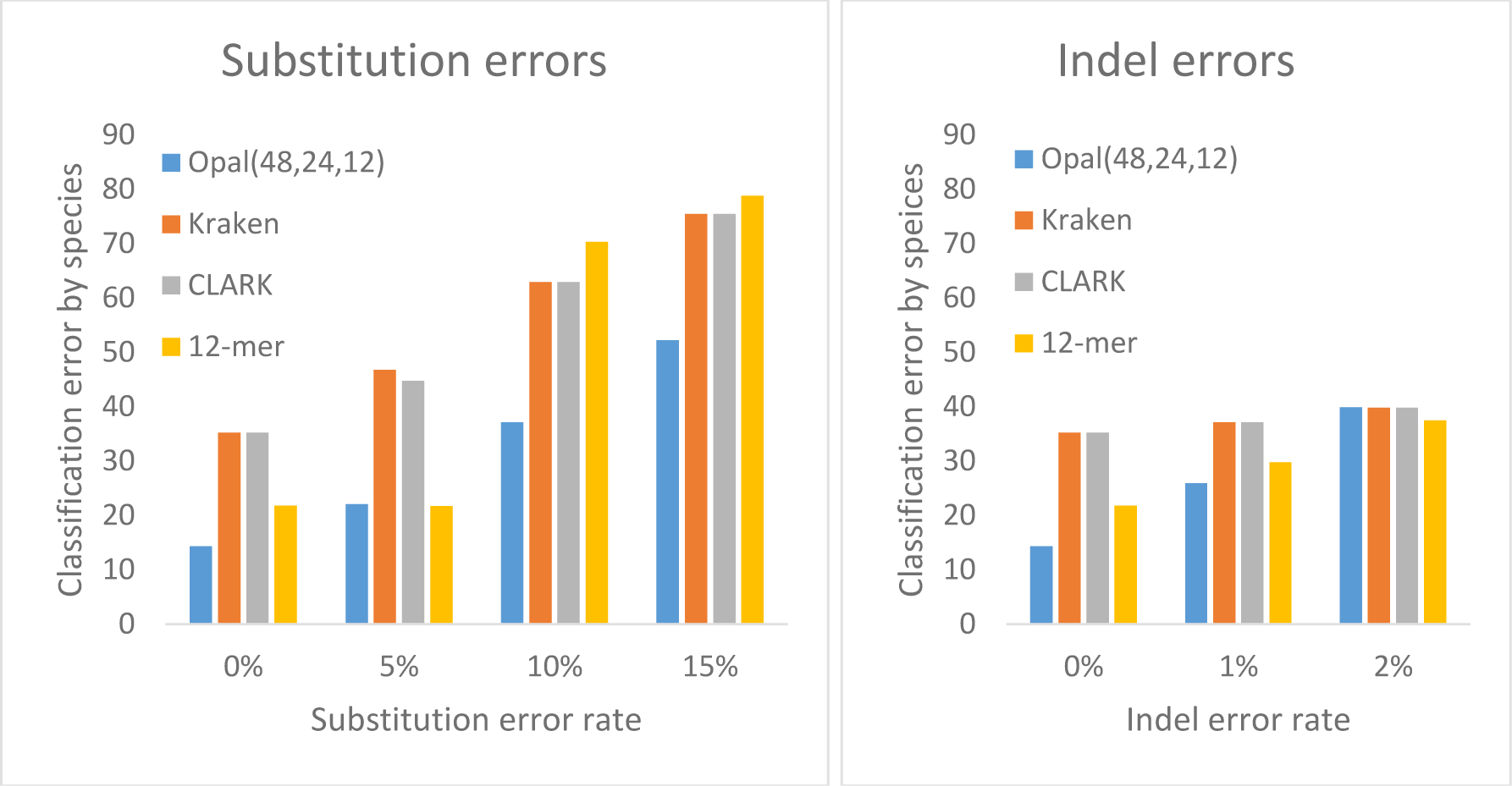
Subspecies benchmark. We compared Opal, Kraken, CLARK, and using contiguous 12-mers on a synthetic benchmark of 7 closely related bacterial substrains with FASTA files from bacteria.ensembl.org:

|  |
| --- |
| Escherichia_coli_o127_h6_str_e2348_69.ASM2654v1.dna.toplevel.fa |
| Escherichia_coli_o139_h28_str_e24377a.ASM1774v1.dna.toplevel.fa |
| Escherichia_coli_o146_h21_str_2010c_3325.Ec2010C-3325.dna.toplevel.fa |
| Escherichia_coli_o157_h7.ASM97884v1.dna.toplevel.fa |
| Escherichia_coli_o7_k1_str_ce10.ASM22762v1.dna.toplevel.fa |
| Escherichia_coli_str_k_12_substr_w3110.ASM1024v1.dna.toplevel.fa |
| Shigella_dysenteriae_1617_gca_000497505.ASM49750v1.dna.toplevel.fa |

Opal was trained with 32 hashes, depth of coverage 820 (4096 batches of 0.2x coverage). The contiguous 12-mer model was trained with depth of coverage 410 (2048 batches of 0.2x coverage). Kraken 0.10.b-beta was run with default options. CLARK v1.2.3 was run with k=31 in full mode. For substitution error rates, as before, Opal performs much better than its competitors. Indel error rate, on the other hand, posed a significant challenge, though Opal still performs comparably at the indel error rates we examined. NB: where Kraken and CLARK chose not to classify a read as one of the 7 substrains given, we count it as randomly guessing, and give 1/7^th^ of a correct classification to its score; this only improves the performance of Kraken and CLARK, providing a more apples-to-apples comparison.

**Table S1.**
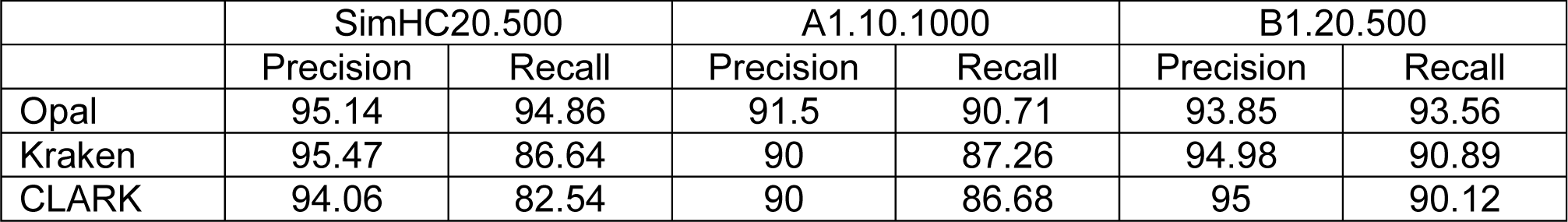
Comparison of Opal against Kraken and CLARK on three benchmarks previously used in the literature. Raw precision and recall numbers for Figure 3A in paper.

|  | SimHC20.500 |  | A1.10.1000 |  | B1.20.500 |  |
| --- | --- | --- | --- | --- | --- | --- |
|  | Precision | Recall | Precision | Recall | Precision | Recall |
| Opal | 95.14 | 94.86 | 91.5 | 90.71 | 93.85 | 93.56 |
| Kraken | 95.47 | 86.64 | 90 | 87.26 | 94.98 | 90.89 |
| CLARK | 94.06 | 82.54 | 90 | 86.68 | 95 | 90.12 |

**Table S2.** Comparison of Opal against Kraken and CLARK at three different phylogenetic levels on a 193-species database previously used in [18]. Raw precision and recall numbers for Figure 3B in paper. Raw sensitivity numbers for novel lineage detection in Figure 3C in paper.

| Binning (193-species large dataset) |  |  |  |  |  |  | Novel Lineage |  |
| --- | --- | --- | --- | --- | --- | --- | --- | --- |
|  | Species |  | Genus |  | Phylum |  | Sensitivity |  |
|  | Precision | Recall | Precision | Recall | Precision | Recall | Genus | Phylum |
| Opal | 86.22 | 76.63 | 90.16 | 89.02 | 95.72 | 95.65 | 31.71 | 22.75 |
| Kraken | 86.1 | 70.31 | 88.33 | 86.08 | 92.29 | 89.83 | 25.35 | 4.85 |
| CLARK | 85.82 | 77.8 | 88.09 | 84.05 | 95.92 | 88.39 | 25.38 | 4.63 |

**Table S3.** Comparison of Opal against Kraken, CLARK, and MetaPhlAn2 at three different phylogenetic levels, measured using Pearson correlation and root mean square error. We calculated Pearson correlation and normalized RMSE between the binning percentages and the actual fractions of reads assigned to their taxonomic origins. Opal clearly outperformed both Kraken and Clark at all levels, likely because Kraken and Clark are based on exact k-mer matches but Opal’s fingerprints can account for mutations or sequencing errors. We also compared Opal against MetaPhlAn2 [30] for metagenomics profiling; even using their (MetaPhlAn2’s) marker genes, Opal performed better on the species and genus level.

| Profiling with all reads |  |  |  |  |  |  |
| --- | --- | --- | --- | --- | --- | --- |
|  | Species |  | Genus |  | Phylum |  |
|  | Pearson | RMSE | Pearson | RMSE | Pearson | RMSE |
| Opal | 0.8154 | 0.6584 | 0.9824 | 0.2262 | 0.9999 | 0.0213 |
| Kraken | 0.8063 | 0.6681 | 0.9612 | 0.2845 | 0.9987 | 0.052 |
| CLARK | 0.7504 | 0.7701 | 0.949 | 0.3266 | 0.9996 | 0.0317 |
| Profiling with all marker genes |  |  |  |  |  |  |
|  | Species |  | Genus |  | Phylum |  |
|  | Pearson | RMSE | Pearson | RMSE | Pearson | RMSE |
| Opal | 0.9695 | 0.2682 | 0.988 | 0.1128 | 0.9999 | 0.0291 |
| MetaPhlAn2 | 0.9541 | 0.3684 | 0.9922 | 0.1622 | 0.9998 | 0.0259 |

